# The MeshCODE to scale – Visualising synaptic binary information

**DOI:** 10.1101/2022.06.16.496395

**Authors:** Samuel F H Barnett, Benjamin T Goult

## Abstract

The Mercator projection map of the world provides a useful, but distorted, view of the relative scale of countries. Current cellular models suffer from a similar distortion. Here, we undertook an in-depth structural analysis of the molecular dimensions in the cell’s computational machinery, the MeshCODE, that is assembled from a meshwork of binary switches in the scaffolding proteins talin and vinculin. Talin contains a series of force-dependent binary switches and each domain switching state introduces quantised step-changes in talin length on a micrometre scale. The average dendritic spine is 1 µm in diameter so this analysis identifies a plausible Gearbox-like mechanism for dynamic regulation of synaptic function, whereby the positioning of enzymes and substrates relative to each other, mechanically-encoded by the MeshCODE switch patterns, might control synaptic transmission. Based on biophysical rules and experimentally derived distances, this analysis yields a novel perspective on biological digital information.

## Introduction

The MeshCODE theory proposes that the meshwork of talin and vinculin molecules scaffolding the synaptic junctions (**Fig.1A**) represent a sophisticated mechanical computation and memory device^1^. Mechanical memory as a concept requires that mechanical signals can be transmitted between neurons with high fidelity and the architecture of the synaptic connections that neurons use to relay signals enable this, as they are discrete, mechanical units isolated from the main cell body. The most common synaptic connection is the dendritic spine and a single neuron can make up to 60,000 of these connections to other neurons to create complex circuits^2^ (**Fig.1A**). The geometry and dimensions of dendritic spines vary but tend to be fairly uniform, composed of a spine head (1 µm diameter) connected to the dendrite by a long spine neck (1 µm in length and 0.1 µm diameter)^3,4^ (**Fig.1A,B**).

**Figure 1:**
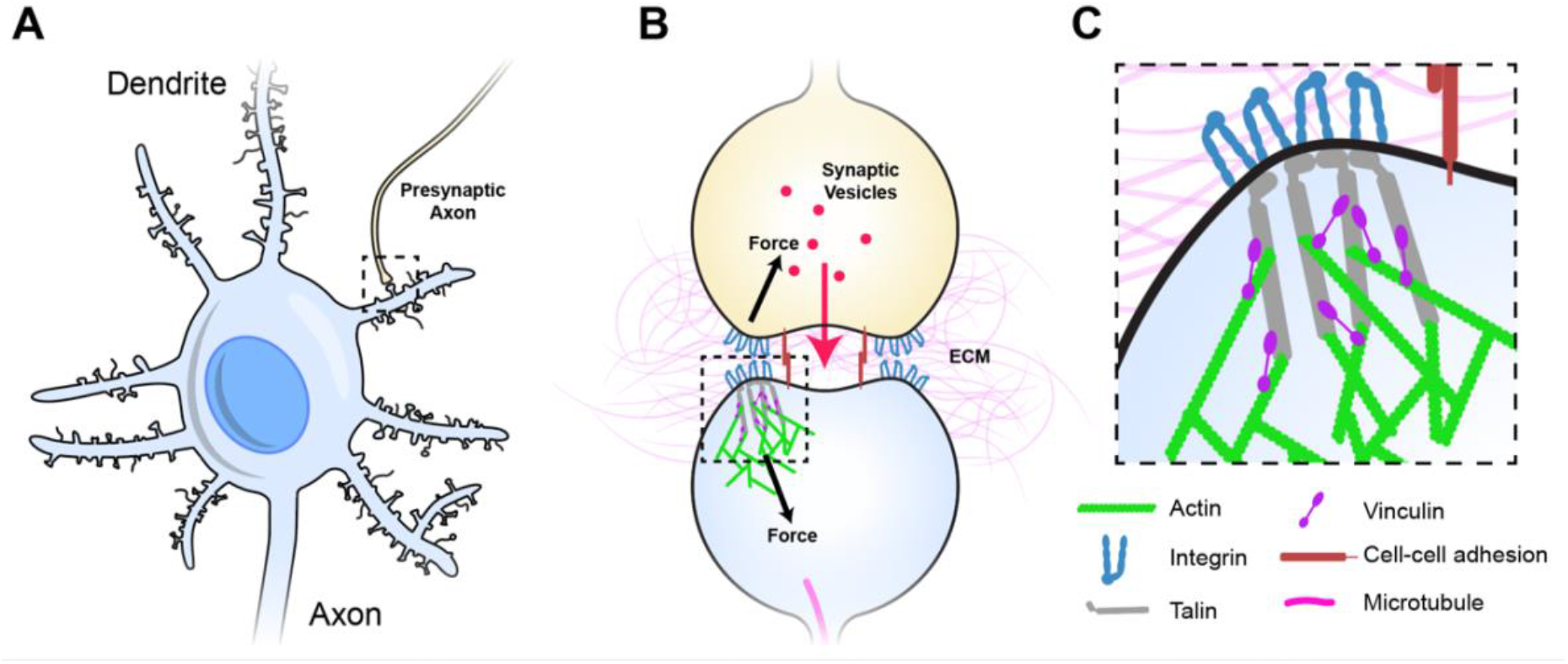
Dendritic spines as mechanical units. **A)** A neuron with multiple dendrites branching from the cell body, each with 100s of dendritic spines (light blue), each of which forms a synapse with an axon of another neuron and receives signals from that neuron. One such connection is shown to an axon from another neuron (yellow). **B)** Typical representation of a dendritic spine showing the pre-(yellow), and post-synaptic (light blue) neurons forming the chemical synapse. The pre- and post-synaptic neurons are mechanically linked, both by sharing the same ECM across the synaptic cleft, but also by cell-cell adhesion complexes (red), these couplings inextricably link the two sides of the junction. **C)** The integrin-talin-actin linkages that define the MeshCODE are shown. Talin (grey) connects the integrin-ECM complexes to the force-generation machinery (green). Vinculin (purple) cross links talin to actin and stabilises the complexes. Figure 1 is not drawn to scale.

Each synapse is surrounded by tightly regulated, mechanically-optimised extracellular matrix (ECM)^5^, and the cells physically engage this ECM forming cell-ECM adhesions around the synaptic periphery that scaffold the synapse^6–9^. The ECM is essential for proper synaptic function and is subject to extensive maintenance^5,10–13^. Integrins are the primary ECM receptors and directly couple the complex meshwork of ECM proteins to the complex meshwork of proteins that is the cells’ cytoskeleton. The ECM-integrin complexes are coupled to the cell’s cytoskeletal machinery via the protein talin. Talin binds the cytoplasmic tails of integrins via an N-terminal FERM domain^14–17^, and mediates cytoskeletal linkages to the actin filaments both directly^18^ and indirectly via the protein vinculin^19^ (**Fig.1B,C**). In this way, talin and vinculin serve to build cytoskeletal structures on the foundations laid out by the integrin:ECM patterns^20,21^. The integrin adhesion complexes (IACs) that assemble on these foundations are essential for neuronal processes including wiring neurons into neural circuits^8^, synaptic plasticity^22^, memory^22,23^ and long term potentiation^24^.

The talin rod region is comprised of 13 switch domains, R1-R13^25^ (**Fig.2A**). These switch domains are helical bundles that open and close in response to mechanical signals^26^. The switches coordinate signalling outputs by recruiting and displacing different signalling molecules as a function of their state^21,27^. When in the open, 1 state, 9 of the 13 talin switches expose binding sites for vinculin^28^. Vinculin stabilises the 1 state of the talin switch it is bound to^29,30^,coordinates signalling via its own helical bundles^31^ and couples to actin^32^ providing additional cytoskeletal linkages and structural reinforcement. The switch patterns and the resultant signalling complexes that assemble provide a mechanism by which information can be carried forward in time where the information can be (re)written by small changes in mechanical force. A signal stimulating synaptic release activates the cells motors specifically at that synapse^33–35^ and in the MeshCODE theory this localised force generation alters the patterns of 1’s and 0’s in the talin molecules on both sides of the stimulated junction and the resultant signalling outputs. The theory proposes that the cells’ entire cytoskeletal architecture serves as a mechanical computation which integrates all the MeshCODEs across the entire cell, with the cytoskeletal linkages to the cell-cell junctions, organelles and the nucleus. Therefore, local changes in the coding at a synapse input into the global mechanical homeostasis computation of the neuron as it resolves all the internal interdependencies within the cell with these updated patterns. When homeostasis is reached, the patterns of 1’s and 0’s across the cell are altered and the entire cytoskeletal machinery updated.

**Figure 2.**
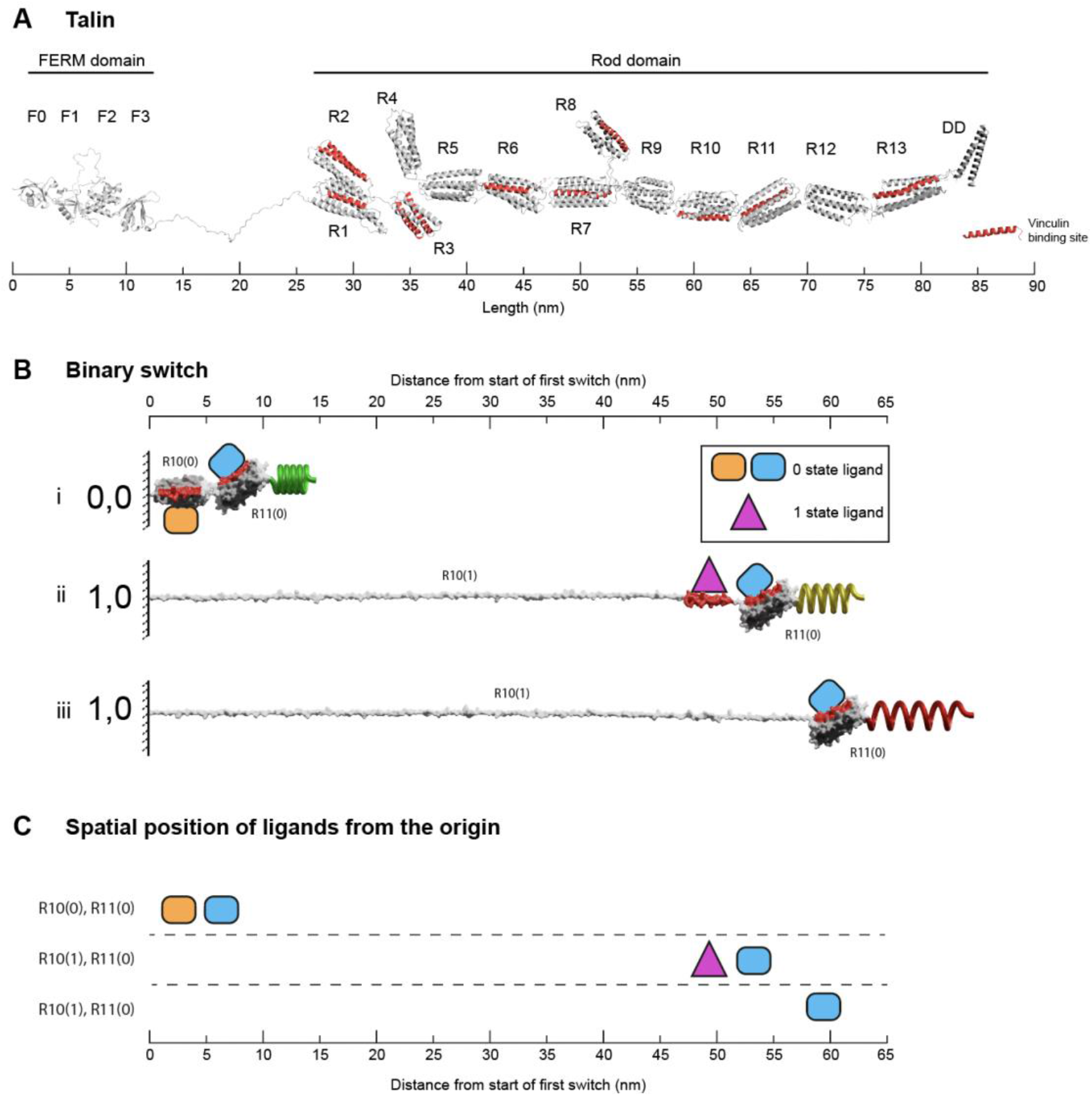
Binary switches in cells. **A)** Structural model of open talin^25^. Talin is composed of an integrin-binding N-terminal FERM domain ‘head’, connected by an 82-residue unstructured linker, to a large flexible rod domain consisting of 61 helices arranged into 13 switch domains (R1–R13). The 62^nd^ helix is a dimerisation domain that forms a dimer with another talin molecule. The 11 VBS helices are shown (red). The length of the talin rod is 55 nm. **B)** The changing dimensions of a talin switch domain *drawn on the same scale as A*. Two 5-helix switches, R10 and R11 are shown. i) In the folded 0,0 state each switch is 5 nm in length. Two signalling molecules are shown bound, orange on R10(0), and blue on R11(0). Both domains contain a vinculin binding site helix (red) that in the 0 state are cryptic and inaccessible. ii) A force increase (shown by a spring) unfolds the first switch to the open, 1 state. After unfolding, the orange R10(0) ligand is displaced and the purple R10(1) ligand engages the now exposed binding site. Vinculin (purple) also limits the switch refolding back to the 0 state. Non-bound helices unravel extending the molecule. Strikingly, the blue ligand remains bound to R11(0), but its location has been altered by 50 nm. iii) Vinculin is displaced, either due to force unfolding the VBS helix or other factors. The fully extended switch is 55 nm, 11x longer than the closed state and comparable in length to the entire fully folded rod in A. **C)** Spatial map of ligands for the two switches in the 0 and 1 states. In this figure the distances are measured from the start of the first switch, however, in the cell these distances would be from the attachment points where the talin is bound to the integrin. All molecules engaging the switches have a coordinate position a fixed distance away from the origin.

Current models of cytoskeletal connections are not drawn to scale which makes it hard to appreciate how altered shapes of the talin “memory” molecules could control synaptic signalling. In this study we performed a detailed structural analysis of the molecules involved in the MeshCODE complexes to build “to-scale” models of the complexes that talin mediates. These scale drawings and videos provide a novel view of IACs and clearly illustrate how the binary switching of talin switch domains can alter the spatial positioning of molecules. Modelling of the mechanically-encoded positioning of molecules within a synapse indicates that each MeshCODE-interacting molecule would have a precise localisation in 3-dimensional space dependent on the switch patterns, resembling a 3-dimensional Venn diagram. We show that by altering the switch patterns, the physical proximity of enzymes relative to their substrates changes, altering the enzymatic activity. The spatial reorganisation of molecules within a sealed unit and primarily in a single dimension resembles the gearbox of a car, and identifies a mechanism for how information in a binary format can control synaptic transmission.

## Results

### The changing dimensions of the force-dependent binary switch domains in talin

Talin has a complex mechanical response due to its 13 force-dependent binary switch domains, so we started this study with a detailed structural analysis of the properties of an individual binary switch as this represents a single bit of information in a digital view of a talin molecule.

There are two types of talin switch: 4-helix and 5-helix bundles. In the absence of force, each switch exists in a compact, folded, 0 state with dimensions 5 × 2 nm, shown drawn to scale in **Fig.2A** (4-helix bundles are slightly narrower in width but the length is 5 nm). A switch will unfold to the open, 1 state in response to tension above its mechanical threshold. In the absence of other factors, this process is reversible and when tension drops it will readily refold back to the 0 state. Opening of a switch sees a transition to an unfolded linear string of alpha helices 1 state. Each helix is ∼5 nm long so the lengths of opened 4-helix and 5-helix bundles are different, being ∼20 nm (4x 5 nm) and ∼25 nm (5x 5 nm), respectively. An exposed talin helix under tension will unravel from the 5 nm helical state to an extended polypeptide form which is ∼12 nm in length. Vinculin binds 11 of the 61 helices in the talin rod^28^ which are located in 9 of the switch domains^25^ (**Fig.2A,B**). Vinculin binds to exposed vinculin-binding sites (VBS) in their helical form, stabilising this conformation^36^ and maintaining it at ∼5 nm in length. As a result, under tension there are two scenarios, exposed helices that do not bind vinculin will extend to 12 nm, but those that bind vinculin will remain at 5 nm, meaning that the length of the switch will be dependent on the ligands engaging it. Vinculin binding also prevents the switch from readily refolding and so stabilises the open state^30^.

Therefore, in the absence of vinculin, a switch opening will transition to a fully extended 1 state. The actual extension will vary, but under tension the contour length from fitting to single-molecule stretching data is ∼0.38 nm/residue^37^. Therefore, starting from the 5 nm folded length, a 4-helix switch extends to 46 nm in length and a 5-helix to 55 nm. That is, a *switch undergoes a 10-fold change in length between the two states* (9-11-fold for 4-helix and 10-12-fold for 5-helix switches). To illustrate this extreme change in the dimensions of a switch, **Fig.2B** shows two 5-helix switches in series. The example domains here are R10 and R11, and the nomenclature used is RXX(0) and RXX(1) to represent the two states. The length of R10 increases from 5 nm for R10(0) to >55 nm for R10(1). In this figure, R11 remains as R11(0) throughout and remains bound to the blue R11(0) ligand. In the R10(0)R11(0) state, both the orange R10(0) and blue R11(0) ligands are located within 10 nm of the origin. Unfolding of R10 displaces the R10(0) ligand that, once released from its fixed location, becomes mobile and able to diffuse away. Simultaneously, the R10(1) ligand binding site is exposed and so the R10(1) ligand which was previously mobile and able to diffuse freely is now restrained, bound to the R10(1) at a fixed position. The R11 switch and its ligand are translocated 50 nm away from the origin. **<u>Supplementary Video 1</u>** shows a movie of another mechanical switching event, here R3 switches binding partners between RIAM (yellow) bound to R3(0) and two vinculins (purple) bound to the R3(1) states as a function of mechanical force^25,38^.

### The changing shape of a talin molecule

The analysis in the previous section illustrates how an individual switch switching states dramatically changes the shape of the domain and the spatial localisation of ligands engaging it. Combining the shape changes of individual switches into a full-length talin molecule with 13 switches, provides a striking visualisation of the dramatic alterations in shape, length and aspect ratio that occur as the switch patterns, and thus the binary information, change (**Fig.3** and **<u>Supplementary Video 2</u> and <u>Supplementary Video 3</u>**). These shape changes alter the signalling molecules recruited and the cytoskeletal connections that emanate from that talin. However, they also result in the relocation of molecules interacting with that talin. Unfolding of a switch domain introduces a step change increase in length of 40 or 50 nm depending on whether the switch that opens is a 4- or 5-helix bundle. R7R8 represents a unique double switch (**Fig.2A**), comprising a 5-helix bundle, R7 with a 4-helix bundle, R8, inserted to form a 9-helix module^39^, opening of this 9-helix module has a step-size of 110 nm (**Fig.3vi,vii**). These step-changes in extension are quantised and a physical property of the talin molecule, since the step size is encoded by the sequence length of the switch. Each switching changes the distance from the integrin attachment site of all molecules engaged to the talin molecule after that switch. Talin-interacting molecules attached either side of a switch will be moved further apart, or closer together, by the size of the step (**Fig.4**). Ultimately, talin can go through many states, a few examples are shown in **Fig.3**: i) autoinhibited (13 nm) ii) integrin-bound but untensioned (86 nm), iii) integrin-bound and tensioned (103 nm), iv) R3(1) (139 nm), v) R2(1)R3(1) (181 nm), all 13 switches in the 1 state (760 nm). The extreme changes in dimension are shown in **<u>Supplementary Video 2</u>**. In migrating cells talin length was shown to fluctuate in the range of 80–350 nm^40^ indicating 0-6 switch domains are unfolded at any one time in dynamic IACs. We note that we are introducing an assumption in these drawings in that the talin molecule is pulled taut. In this analysis we are looking at the end-to-end distance as defining the maximum extension of the mesh.

**Figure 3.**
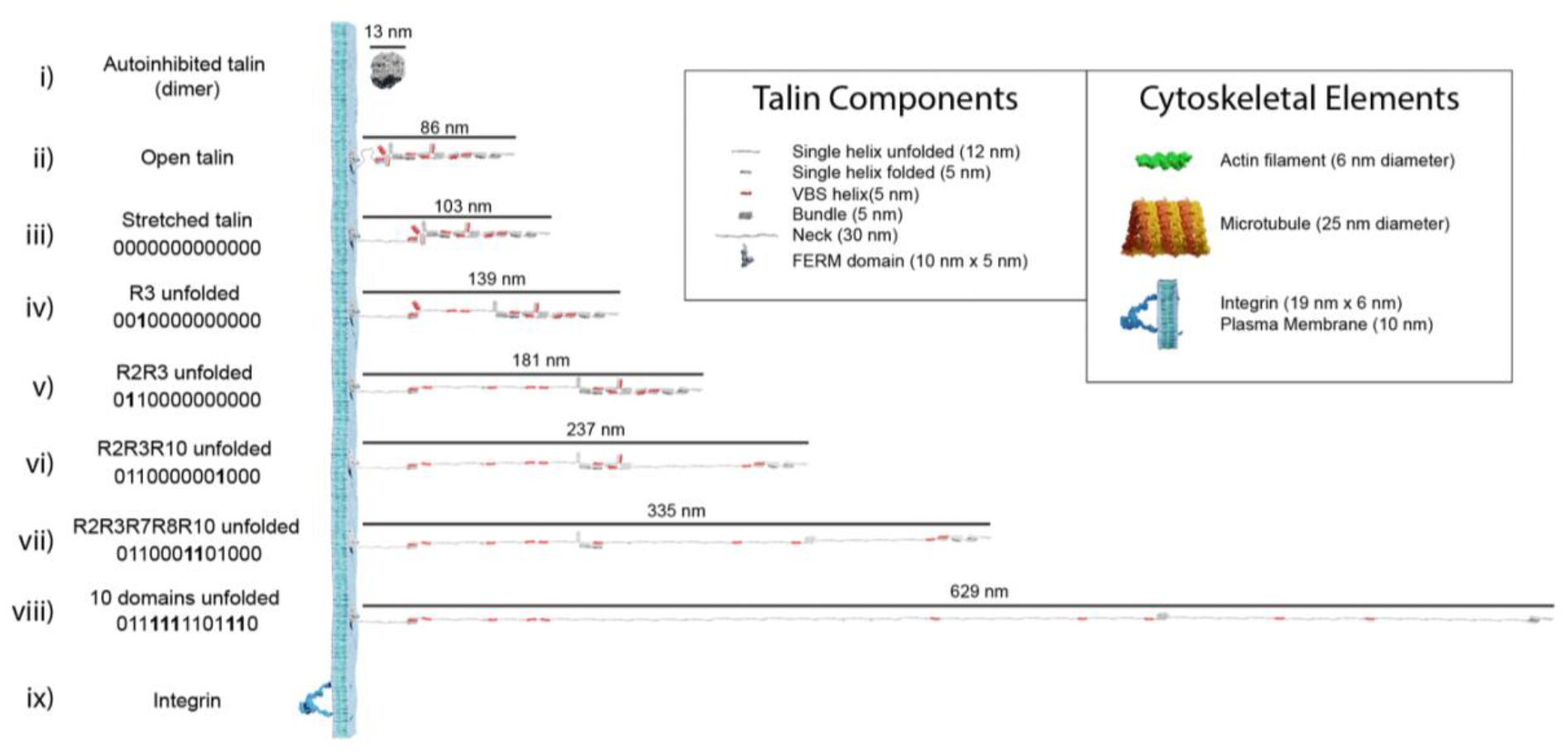
Talin to scale. Scale models of talin in different conformations. The length of each form is shown in nanometres. The binary string represented by each state is shown, with 0000000000000 representing the fully folded state and 1111111111111 the fully open state. In each subsequent binary string, the changing switch is indicated in bold. i-iii) Three representations of fully 0-state talin, i) The autoinhibited dimeric form of talin^41^, the autoinhibited monomeric form^42^ is slightly smaller. ii) Open talin, the same as in Figure 2A is ∼86 nm in length. iii) As talin experiences tension, the linker can extend so the molecule is ∼103 nm. iv) As force is exerted on talin, the R3 switch opens first, exposing two VBS, and the length of the molecule increases by 36 nm if both VBS bind vinculin, or 47 nm otherwise. v-viii) As additional domains unfold, the binary patterns change and the length of the molecule undergoes stepwise changes in length, ranging from 86–760 nm whilst still maintaining a mechanical linkage. Only 10 domains are shown unfolded in viii) but 12 can unfold and still maintain the mechanical linkage to actin via R13. We note that it is possible for all 13 switches to unfold and still maintain a mechanical linkage at the C-termini via the VBS in R13. The fully 1-state talin is not shown as its length would exceed the page width. ix) Remarkably, integrins are relatively small in comparison, here an active integrin heterodimer is shown on the same scale. Inset: The dimensions of the different talin components. Major cytoskeletal elements including integrin/plasma membrane, actin filaments and microtubules are shown for comparison. The integrin cytoplasmic tails which talin connects to extend ∼10-15 nm.

### The MeshCODE data patterns have vastly different geometries and dimensions

Putting the dimensions of the individual molecules together, it is now possible to visualise the impacts that alterations to the MeshCODE switch patterns will have on the synapse. A synaptic signalling event would lead to the motors being switched on inside that synapse^35^ and the resultant pulling on the MeshCODE would alter the shapes of the talin switches^43^. As shown in **Fig.4**, a single switch unfolding leads to significant alterations in the sub-cellular localisations of mesh-interacting proteins. These alterations in spatial positioning occur along a straight line between the origin, which can be considered the tether point where the talin engages the ECM:integrin at the plasma membrane, and the site of force loading on the molecule. As a result, molecules engaging talin are spatially positioned in the z-dimension and will be relocated along that axis in quantised steps as a function of the changing switch patterns.

**Figure 4.**
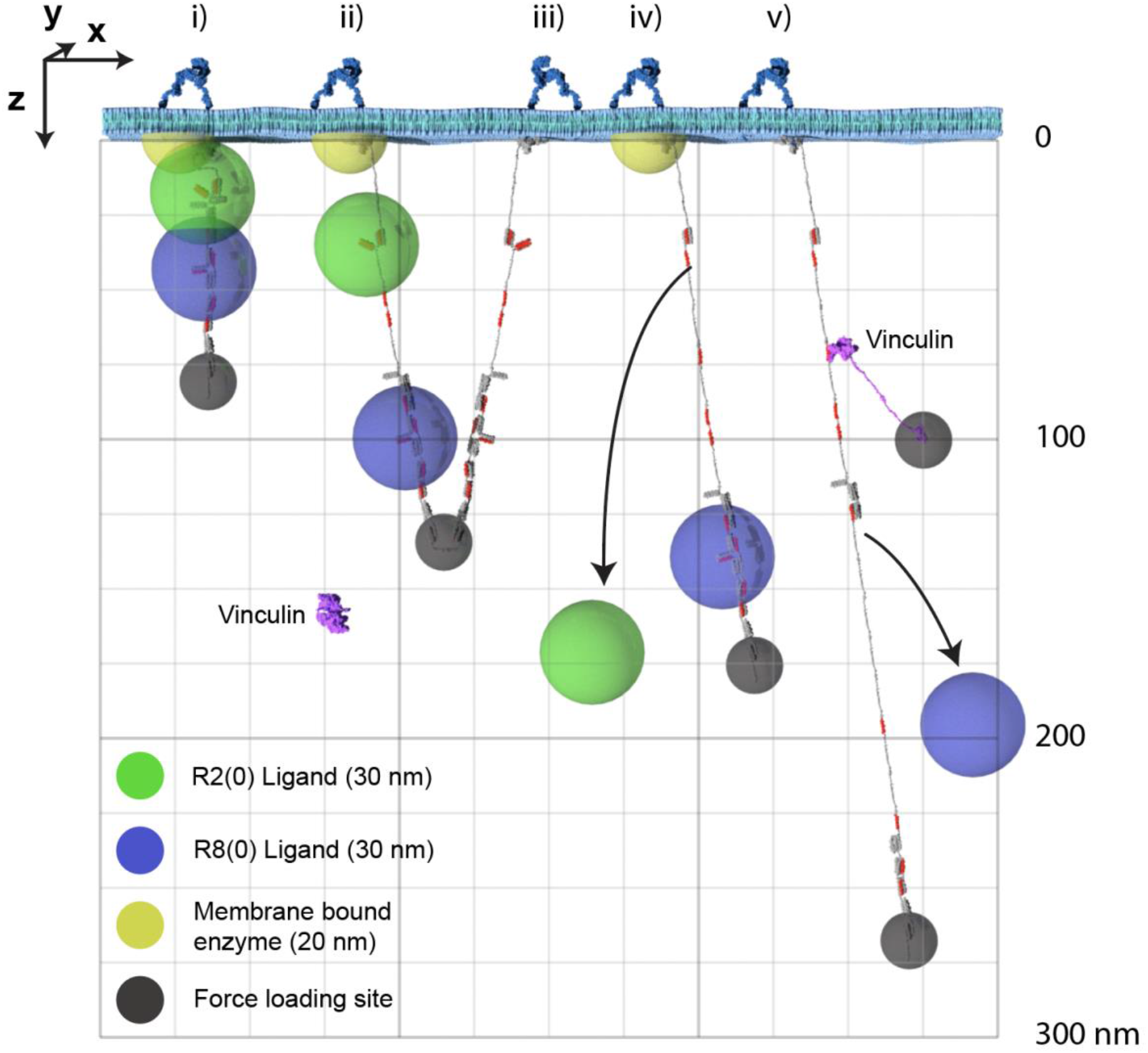
The MeshCODE defines a 3D Venn diagram of molecules spatially resolved in the z-dimension. The x- and y-dimensions are the plane of the membrane with z-being the axis away from the membrane. For clarity only two dimensions, x- and z- are shown. Five talin molecules are shown drawn to scale and in different states (i-v). The yellow hemisphere is a membrane tethered enzyme or substrate. The green/blue spheres represent ligands engaging the MeshCODE and illustrate the zones of activity of molecules engaging the switches. The green sphere is an R2(0) ligand, the blue sphere is an R8(0) ligand, and the small dark grey spheres represent the mechanical attachments exerting the pulling force in the z-dimension. A single vinculin is drawn, autoinhibited in the cytosol and then bound to a VBS in R2(1). Each molecule has a fixed location which can be given Cartesian x,y,z coordinates relative to the fixed origin position defined by the ECM-integrin-talin connection. In some switch states these spheres overlap, but as the switch patterns change, the positions of each sphere are changed relative to each other. For clarity we only show the result of 3 switches opening as additional switches opening would exceed the page height at this scale. Many talin ligands are enzymes or known enzyme substrates, so where these spheres overlap the enzymatic activity will be high, and where they are held apart the activity will be low. The size of the spheres where each ligand is active will be dictated by the properties of the ligand but the locations of the spheres relative to each other will be a function of the switch patterns.

### Enzymatic activity in the synapse is spatially restrained

The spatial organisation of enzymatic activity can be achieved via scaffold, anchoring and adapter proteins^50^. Many talin ligands are enzymes, or known enzyme substrates^27,44,45^, and there are many enzymes linked to IACs^46^ and located in dendritic spines^47^. Some of these, such as receptor-linked kinases are membrane bound, some are freely diffusing and some specifically engage the talin switches. Focal Adhesion Kinase (FAK)^48^ and Cyclin Dependent Kinase 1 (CDK1)^44^ have both been shown to bind to talin. Recently, the acetyltransferase, alpha-tubulin acetyl transferase-1 (ATAT-1) has also been reported to directly bind talin to mediate its effects on microtubule dynamics^49^. ATAT-1 and FAK have large unstructured linkers between their anchoring points and their enzymatic domains (**Fig.5**). If an enzyme has an unstructured region between the enzymatic domain and the anchor site, then that unstructured linker defines a “zone of activity” for that enzyme (**Fig.4,5** and **Supplementary Movie 4**). As an analogy, one can consider a goat tethered to a stake in a grassy field. The goat will eat all the grass in the region it can access, which is dictated by the length of its tether and the position of the stake. Grass outside of this zone of activity will not be eaten. Repositioning the stake will enable the goat to access different regions of grass, and altering the length of the tether will dictate the size of the circular patch of eaten grass it will make. Similarly, a kinase can phosphorylate any substrate present within its zone of activity, which we draw as a sphere with a radius of the linker length. If tethered, these zones of activity are positioned in precisely defined spatial locations within the cell, able to reach substrates efficiently only within that sphere. If the anchor site is to a talin switch domain, then these spheres are positioned at specific distances away from the integrin attachment point dependent on the switch pattern of that talin. In such a scenario, a talin switch unfolding between an enzyme anchor and the membrane will move the enzymes’ zone of activity a quantised distance away from the membrane. Alternatively, a switch unfolding might move meshwork-bound substrates out of reach of membrane-bound enzymes. In this way, tight control of signal transduction pathways can be maintained via the spatial positioning of enzymes with opposing functions (such as kinases and phosphatases), relative to each other and their substrates, mechanically-encoded by the MeshCODE switch patterns. Transiently increasing the amount, activity, or localisation of one enzyme relative to the other is a way to ensure specificity of signal transduction at that site.

**Figure 5.**
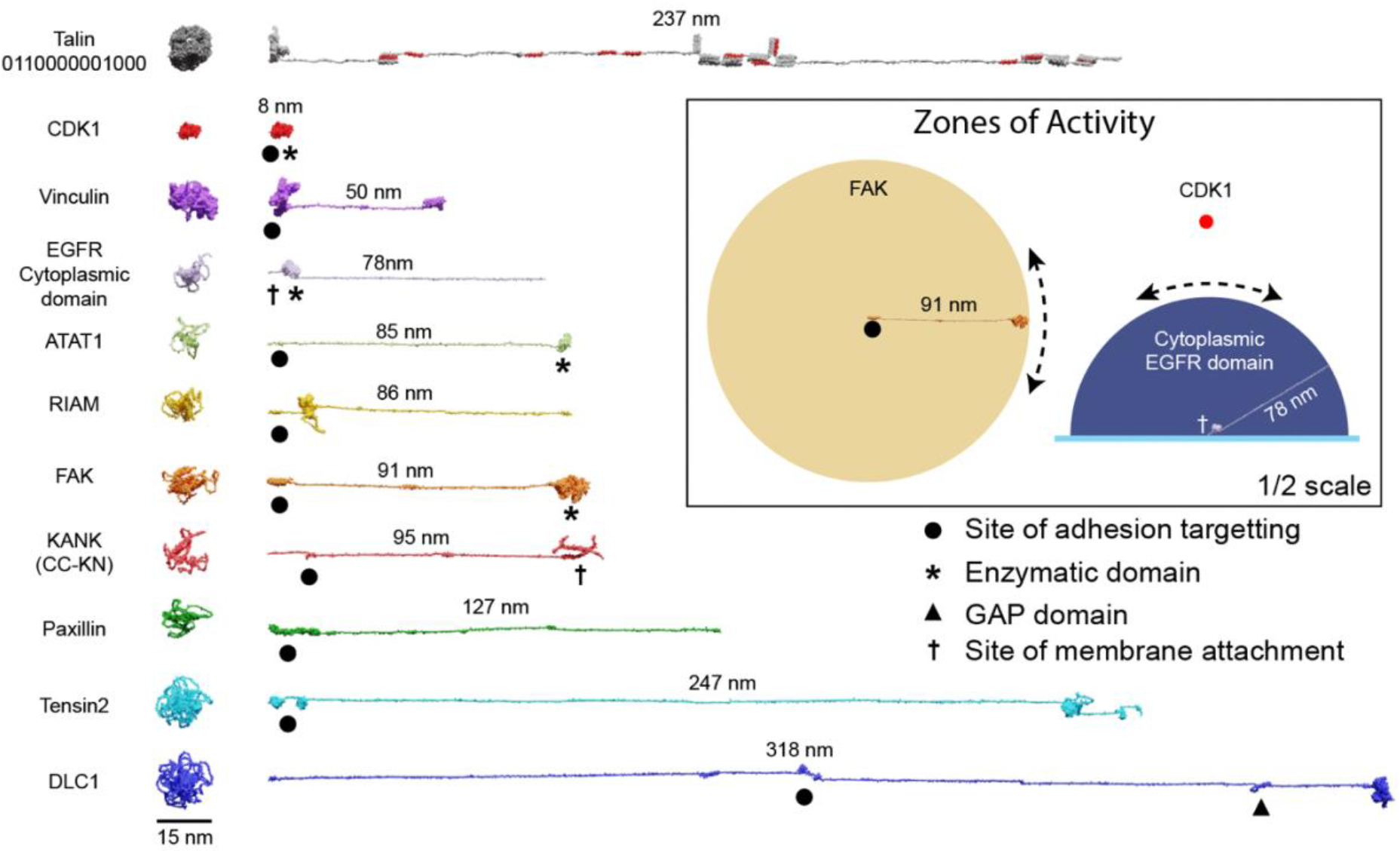
Zones of activity of MeshCODE interacting proteins are dictated by their unstructured linker regions. AlphaFold models of a subset of the 250 components of IACs drawn to scale. The compact AlphaFold model (left) and extended version (right) are shown for each to illustrate the length of these proteins. Talin is included as a scale bar. The length of the unstructured region defines the radius of the sphere of activity of that protein; when attached to the MeshCODE it can reach molecules within that sphere. Membrane-tethered enzymes have a hemisphere zone of activity, Epidermal Growth Factor Receptor (EGFR) is shown as an example of a membrane-tethered receptor tyrosine kinase (only the cytoplasmic region is shown). Enzymatic domains are marked by * and the GTPase-activating protein (GAP) domain by a triangle. Focal adhesion targeting regions are shown by a dot and membrane attachment sites by a cross. For KANK only the region from the membrane-targeting coiled coil to the talin binding site at the N-terminus is shown (residues 1-517). **Inset**: Zones of activity of three MeshCODE interacting enzymes, Focal Adhesion Kinase (FAK), Cyclin Dependent Kinase 1 (CDK1) and the membrane-tethered EGFR. The radius of the sphere is the length of the linker between the anchor point and the enzymatic domain, in the case of FAK this is 91 nm, for CDK1, the enzymatic domain binds directly to talin so the sphere is much tighter. The EGFR hemisphere has radius 78 nm. The location of the sphere in 3D space is dictated by where the molecule is anchored, if the anchor point is to the membrane it will be relatively constant, but if it engages talin either directly or indirectly, its location will change dictated by the switch patterns as shown in **Fig.4**. For membrane tethered enzymes the position in z-will be constant. The inset is shown at half scale compared to the rest of the figure.

### The MeshCODE to scale inside of a dendritic spine

The architecture of a dendritic spine consists of a bulbous head connected to the main dendrite via a spine neck region (**Fig.1** and **6**). Drawn to the same scale as talin it becomes possible to visualise what binary information in the MeshCODE of a dendritic spine would look like (**Fig.6**). The integrins are positioned around the active zone of the synapse and the force-generation machinery acts from the bottom of the spine. Each time a switch domain unfolds, it results in a quantised extension of the talin molecule, predominantly in a single dimension defined by a straight line from the integrin attachment site towards the bottom of the spine (**Fig.6A,B**). This indicates that the binary patterns will dynamically alter the spatial positioning of molecules interacting with the talin switches and the cytoskeletal linkages emanating from them. A protein bound towards the C-terminus of talin, or to a downstream cytoskeletal connection, could see itself relocate >800 nm in the z-dimension following unfolding events, whilst remaining physically coupled to the active zone/PSD. This spatial reorganisation of the same molecules bound to the same integrin-talin complexes is shown in **Fig.6C,D** where four talins are illustrated bound to ligands, each with a zone of activity (**Fig.6E**). In **Fig.6C** the talin switches are mostly closed and the bound molecules are concentrated at the active zone/PSD. In contrast, in **Fig.6D** where more talin switches are opened the same molecules are organised differently, with greater separation between them in 3D-space. In **Fig.6D** a role for vinculin (purple circles) is apparent, helping maintain the distance between the bound ligands.

**Figure 6.**
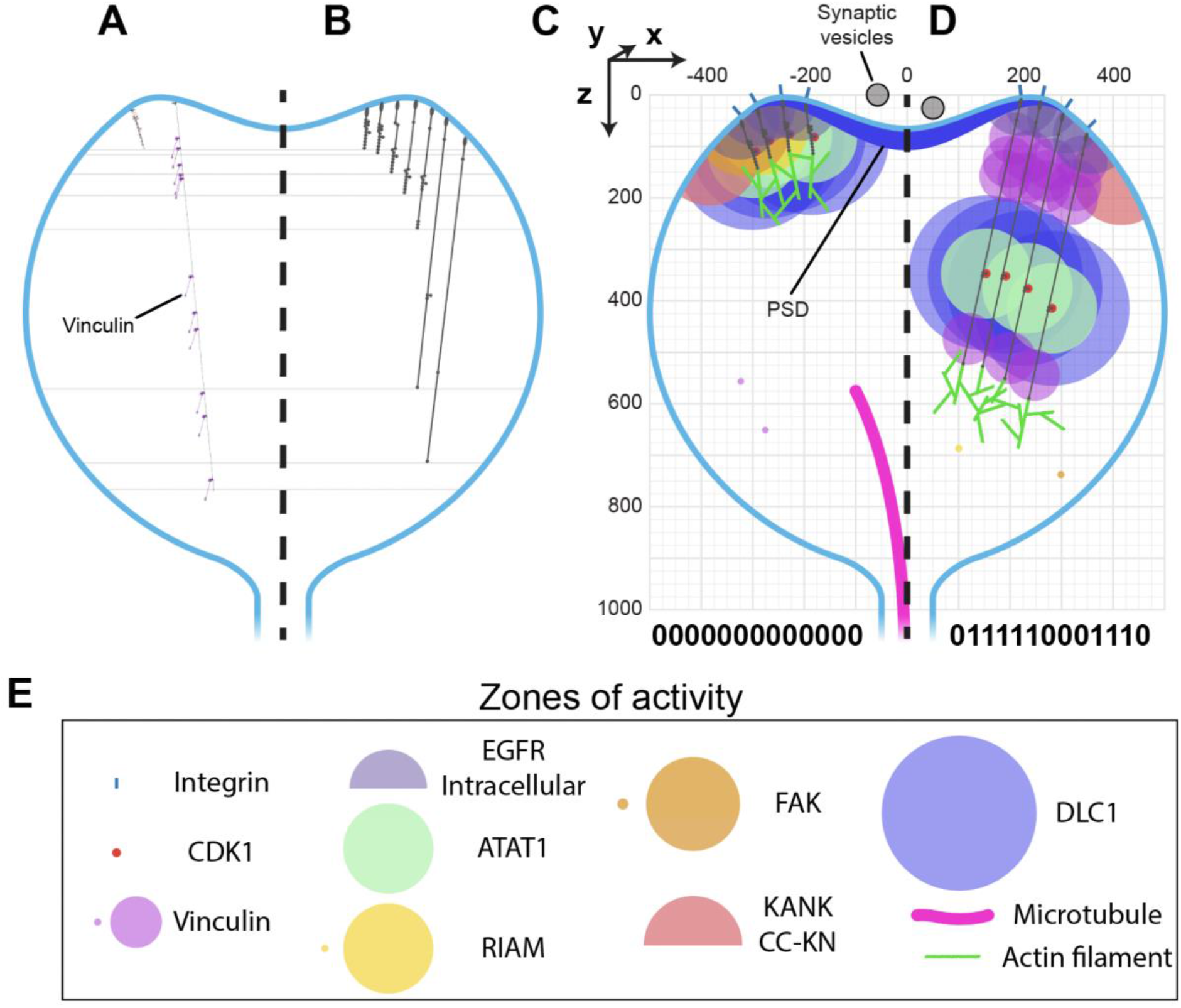
MeshCODE to scale inside a dendritic spine indicates a mechanism for mechanical coordination of synaptic activity. **A-D)** Scale drawings of a dendritic spine (1 µm diameter), with talin in different switch states. **A)** The talin extremes, the fully folded 0-state (left) is 86 nm and the fully extended 1-state (right) is 790 nm. The spatial positioning of the 11 VBS is shown with 11 vinculin molecules (purple) engaged. **B)** Different talin states define layers in the z-dimension where binding sites are positioned. Molecules engaging talin will be spatially positioned between 0-700 nm from the integrin attachment site depending on the switch pattern. **C-E)** Scale drawing of a spine showing the zones of activity of talin interacting molecules. A three-dimensional coordinate system is used, with the origin positioned at the centre of the active zone/PSD. This allows each molecule’s location to be defined with a Cartesian x,y,z coordinate, and distances between them described mathematically. The quantised steps in talin length move engaged spheres a precise distance in the direction of the force vector, which is a straight line away from the integrin attachment site around the active zone/PSD towards the spine neck. In this way the binary coding of the synapse will organise molecules at fixed positions relative to the origin. Four integrin-bound talin molecules are shown in **C)** compact, and **D)** extended states highlighting how the engaged molecules spatially resolve into layers as a function of the binary switch patterns. **E)** Zones of activity of the molecules in C-D. Where two zones are shown, the smaller of the two represents the autoinhibited state and the larger the active state.

The spatial organisation of molecules in the synapse will be coordinated by the binary patterns in the talin molecules which can be dynamically altered by synaptic signalling. Not all molecules will directly engage the MeshCODE, many synaptic proteins will be anchored at the active zone/PSD and so will not change position when the switch patterns change. This indicates a mechanism for how binary information could control synaptic activity whereby the spatial reorganisation of enzymes and substrates as the binary strings change would dynamically alter the rates of reactions.

### The Synaptic Gearbox Model - A novel hypothesis for the dynamic regulation of synaptic activity via binary MeshCODE patterns

Visualising the MeshCODE within the confines of a synapse identifies a novel mechanism for how synaptic activity could be dynamically regulated. Mechanical reorganisation of molecules within a spine would alter the activity and efficiency of the enzymatic reactions and thus the signalling at that spine (**Fig.6**). The talin molecules anchored around the periphery of the active zone/PSD, would be acting in synchrony as springs, connecting the force generation machinery to the active zone. Each talin would be serving as a mechanosensitive signalling hub^21,27^ and as the binary patterns change, molecules engaged to the MeshCODE would be (re)positioned at quantised distances relative to the active zone/PSD. If all the talins within a spine have similar binary patterns it would have the implication that molecules engaging the MeshCODE would be stratified in the z-dimension relative to the active zone/PSD (**Fig.6C-D**). In contrast, molecules not engaged to the MeshCODE would either stay where they were if tethered to the active zone/PSD (or elsewhere) or be freely diffusing. This identifies two ordered components of the synaptic machinery, the “static component” that stays near the active zone/PSD, and the “mechanically operated, MeshCODE-interacting component” that can move closer and further away from the active zone as a function of the switch patterns (**Fig.6F-G**). Enzymatic reactions have a strong concentration dependence, so spatial separation of enzyme from substrate would dramatically alter the local concentrations and the efficiency of the reactions and thus the signalling at that spine (**Fig.4,6**).

In this way the MeshCODE would coordinate synaptic activity by moving the two parts of the machinery relative to each other, in a graduated fashion, almost like the gearbox on a car (**Fig.7**). By moving the mechanically operated part of the synaptic machinery relative to the active zone/PSD the activity of the synapse can be tightly controlled and modulated by quantised step-changes in distance. The morphology of the synapse supports such a mechanism as this mechanical tuning of synapse activity would occur primarily within the bulbous head, discrete from the rest of the neuron. The thin, yet long, connecting neck of the spine would mechanically separate the synapse, and also limit free diffusion into and out of the spine, serving as a kinetic trap for molecules. In such an apparatus the exposure, closure and relocation of binding sites within the confines of the synapse will have outsized effects on local molecular processes.

**Figure 7.**
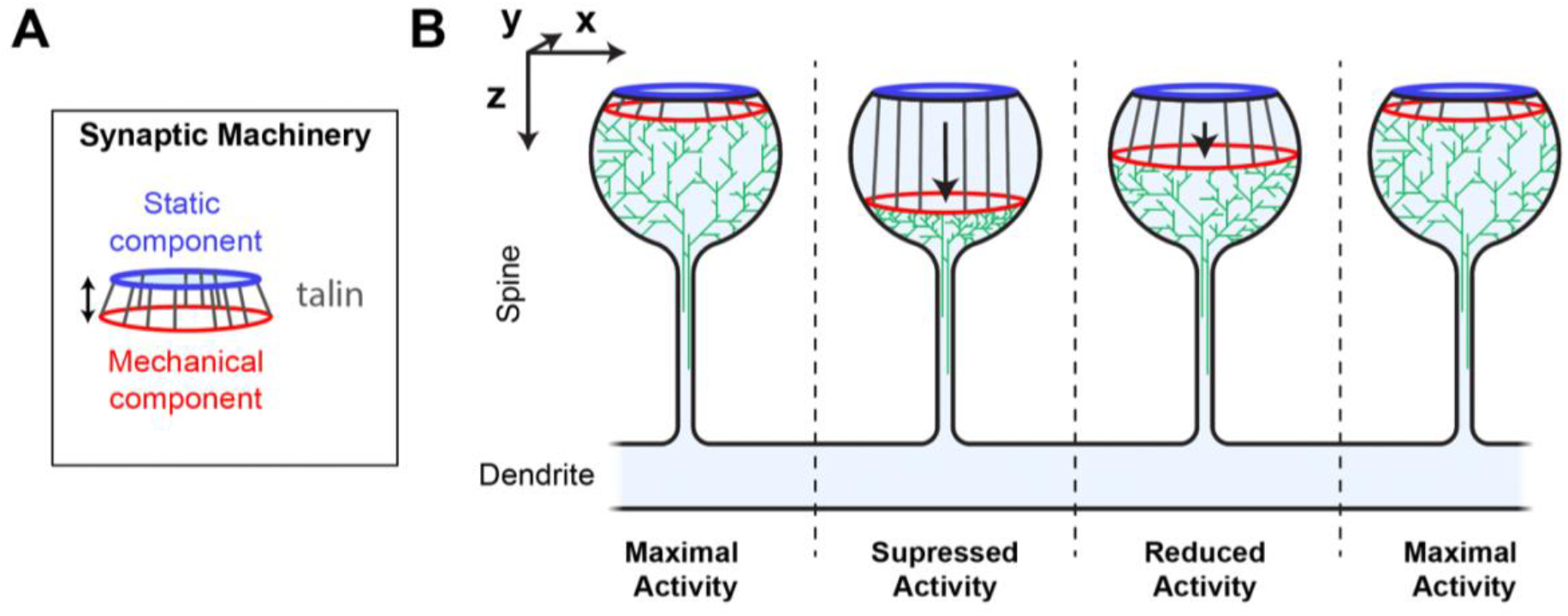
Synaptic Gearbox hypothesis of a dynamically tuneable, binary-encoded mechanism for graduated synaptic activity. **A)** The synaptic machinery can be divided into two ordered parts, i) a static component at the active zone/PSD (blue) and a mechanically operated component (red) connected by the MeshCODE (grey). The graduated nature of the MeshCODE switch states means that synaptic activity can be dynamically regulated by moving the two parts of the machinery relative to each other in quantised steps. **B)** Despite having similar molecular compositions, the four synapses illustrated are shown as having different activation states because the molecules in the synapse are ordered differently due to different MeshCODE patterns. Mechanical signals that change the MeshCODE patterns will re-organise the synaptic machinery and dynamically tune up or down the synaptic activity. In this way every synapse in the organism can be mechanically synchronised by altering the binary coding within each.

### Global and persistent restructuring of the MeshCODE by posttranslational modifications to the mesh

Our recent study of the direct interaction between talin and CDK1^44^ identified an intimate coupling between mechanical and biochemical signalling, hinting at how the MeshCODE and enzymes might interact to give diverse outcomes. CDK1 binds directly to R8(0), leading to phosphorylation of multiple IAC proteins^51^, including talin itself on R7 at serine1589^44^. The R7(0)R8(0) state has similar length to the other switches, with a 5 nm branch where R8 protrudes from the talin rod (**Fig.2A**). However, upon switching states R7R8 undergoes a 22x increase in length, from 5 to 112 nm. As a result, major cytoskeletal and compositional changes occur when R7 switches state. Phosphorylation of Ser1589 alters the mechanical response of R7R8, lowering the unfolding-force threshold from >14 to ∼10 pN^44^. As illustrated in **Fig.3vi-vii** and **<u>Supplementary Movie 5</u>** this posttranslational modification (PTM) dramatically alters the MeshCODE dimensions and cytoskeletal connections. The R7R8 switches have emerged as a major nexus of cytoskeletal dynamics coupling the actin^19,52^, and microtubule^53^ cytoskeletons. R7(0) coordinates the targeting of microtubules to IACs via a direct interaction with the KANK family of proteins^53^ and R8(0) forms part of a second actin-binding site, ABS2^19,52^. Furthermore, R8(0) is a binding site for numerous LD motif-containing ligands including CDK1^44^, RIAM^25^, DLC1^54^, paxillin^54^ and α-synemin^55^. Therefore, R7R8 unfolding has 6 major impacts on the MeshCODE, as it: i) causes >107 nm increase in talin length, ii) disrupts actin binding to ABS2 iii) displaces LD motif-containing ligands from R8(0) iv) disrupts microtubule targeting to IACs, v) exposes two VBS, in R7(1) and R8(1) and vi) moves ligands for R9-R13 and downstream attachments, 110 nm away from the origin.

These extensive changes, all resulting from a single PTM of a binary switch, indicate a mechanism for how chemical and mechanical signals can integrate to coordinate the 3-dimensional organisation of molecules to control cell functioning. There are numerous post-translation modifications on talin switches^27^ which likely integrate chemical and mechanical signalling into precise MeshCODE patterns that help fix the 3-dimensional locations of molecules within the cell.

## Discussion

The concept of a cellular binary coding coordinating organisms is currently contentious. However, experimentally it is well shown that each talin molecule is built of a string of binary switches^26^, able to coordinate different cellular processes as a function of the patterns of folded, 0 and unfolded, 1 switch domains^21,27^. Whilst it is established that talin is a memory molecule, and IACs are a central component of the synaptic scaffolding organising each synapse^6,8^, it is not yet demonstrated that the memory aspect of talin is acting in this context. In this work we sought to use the wealth of structural information about talin and its binding partners to build experimentally derived models of the complex meshwork of binary switches, the MeshCODE, that assembles on the cytoplasmic face of integrins around the edge of each synapse.

Our analysis indicates that the spatial organisation of molecules within a cell might be extensive, with proteins organised up to 800 nm apart whilst still physically coupled together. The length of talin and the step changes that result from alterations in its binary coding are highly conserved across evolutionary time (**Fig.S2**) supporting its role as a dynamic, spatial organiser of molecules. This leads us to propose that IACs are likely to be meticulously ordered in all 3-dimensions *in vivo* with the >250 different proteins involved in these complexes^46^ spatially organised relative to each other. Vastly more information and complexity would be possible in such an ordered system. Counterintuitively, higher ordering of molecules necessitates larger disordered regions in the proteins themselves because, as proteins are restrained to certain locations within a cell, unstructured regions of proteins enable them to function within discrete regions which we term zones of activity (**Fig.4-6**). Approximately 30% of the human proteome is predicted to be disordered, and unstructured protein regions are more prevalent in complex multicellular organisms than in simple unicellular organisms^56^ consistent with greater spatial organisation of molecules in more complex systems. Many researchers who have looked at their protein of interest using the protein structure prediction software, AlphaFold^57^ will have observed the amount of “spaghetti-like” unstructured regions present in many proteins (**Fig.5** and **S1**), contrasting starkly with the ordered atomic structures of the folded domains. As tethered linkers, the key aspect of these unstructured regions would be the length it can extend, enabling proteins to engage their targets within discrete regions of the cell. Therefore, conservation of the optimal linker length might be more important than the sequence itself. Another observation we make is that MeshCODE ligands fall into two distinct categories, proteins with large linker regions which will have zones of activity and/or engage multiple nodes on the meshwork, and globular proteins that likely diffuse in and around the meshwork and attach as and when binding sites become available to exert more localised effects.

This ordering of molecules as a function of the switch patterns will organise the post-synaptic force generation machinery (**Fig.6,7**), which will alter the ECM and cell-cell linkages, modulating the effective stiffness the pre-synaptic neuron experiences. As rigidity sensing is an established phenomenon that occurs via IACs^58,59^ a stiffer interface presented to the pre-synaptic neuron will alter its mechanosensing and so synaptic vesicular release could be dependent on the post-synaptic side via a mechanical mechanism where the binary patterns in both the pre- and post-synapse contribute to the overall activity of the synapse. Each synaptic release would alter the coding and the activity of that synapse in relation to future inputs. It might be possible to determine the probability of vesicular release by measuring the current effective stiffness of the synapse, or the relative positioning of the static and mechanically-operated parts of the machinery. The status of each synapse would feed into the global mechanical computation of each and every neuron and so each signalling event would update this global coding. In this way, every neuron in the organism would become mechanically synchronised, with a complex binary code operating throughout.

Francis Crick first raised the issue that biomolecular information storage is limited by molecular turnover^60^. Information written in the shapes of molecules faces an additional issue that molecules are dynamic and thermal noise means no system is perfect. The switch patterns by themselves are stable under low tension for minutes to hours^26^, however, proteins like vinculin additionally stabilise these patterns^30,61^. Furthermore, residues that are buried in the switches have been found to be phosphorylated in proteomics screens^62–64^. Such residues would be exposed only when the switch was open, and so phosphorylation would limit refolding, fixing the switch in the 1 state. This indicates that the patterns themselves can be long-lived and the large complexes that form on the meshwork would convert these patterns into biological outputs. However, the molecules themselves would still be subject to the natural turnover of biomolecules - although tethered molecules held in place under tension would be subject to reduced turnover. As part of the mechanically-operated machinery all the talin molecules would be working together to spatially organise a significant part of the synaptic apparatus, and a single talin molecule detaching would not suddenly break this mechanism, as it could be compensated by the rest of the machinery. When a talin molecule does untether and gets turned over, then the spatial patterning would be perturbed which would necessitate rebalancing to re-establish mechanical homeostasis. This rebalancing would result in the synapse replacing that talin molecule and rewriting the molecular shapes *provided that they are consistent with the rest of the mechanical computation*. If the information was inconsistent, then this recalculation would necessitate larger changes to re-establish mechanical homeostasis. In this way the turnover of molecules need not limit biomolecular information storage in the shape of molecules and may even help to ensure the whole neuron, and by extension the whole neuronal circuitry, remains mechanically synchronised.

In summary, visualising molecules to scale has identified a previously unrecognised role of mechanotransduction whereby, alterations in the binary patterns in the shape of the MeshCODE protein talin lead to the spatial reorganisation of molecules. This reorganisation, occurring primarily in a single dimension away from the attachment site, indicates a novel Gearbox hypothesis for how mechanical switches might coordinate synaptic transmission. Information written into the synaptic scaffolds in binary format would result in the precise spatial positioning of molecules within the synapse, with each switching event dynamically altering the distance between enzymes and their substrates and in doing so their activity. A mathematical representation of the synaptic architecture, with the locations of each molecule represented by Cartesian coordinates that reposition by quantised distances in response to changes in binary coding might offer a more quantitative way to consider synaptic functioning. Beyond the synapse, our scale drawings also offer a deeper appreciation of the architecture of what the inner workings of other cell types might look like, where the dynamic reorganisation of molecules would provide the capacity for much greater cellular information content than current models allow. The videos we present of a talin altering its shape as the switches open and close in response to force illustrate the incredible amount of information that can be written into the shape of each talin molecule. Finally, these representations of memory molecules provide a visual metaphor for binary cellular information with far-reaching implications for how information-processing in animals might be occurring.

## Methods

### Structural analysis

To determine the size of each protein domain we used the atomic structures obtained from the Protein Data Bank (PDB) files. Structural models collected from the EMBL AlphaFold2 database^57,65^ were used for the visualisation of unstructured regions. This was particularly useful for the proteins where only the folded globular regions have structural information available. Domain measurements were taken as the major/minor dimensions as assessed in PyMOL (Version 2.0.5 Schrödinger, LLC), and the “Measure Distance” plugin was used to get accurate sizes. The Aspect Ratio (AR) for the talin rod bundles was calculated using the ratio of the major/minor dimensions. As the rod domains are almost cylindrical, the minor dimension represents the cross-sectional width of the domain. For the creation of extended proteins, the AlphaFold model each amino acid was serially rotated in PyMOL to “linearise” the unstructured elements. Domain boundaries were identified using the AlphaFold2 structures.

### Animations

For the construction of the talin model used in the animations, the crystal structure of the FERM domain (PDB 6VGU^66^) was fused to an 82 amino acid stretch of unstructured linker from the Tensin AlphaFold model. For the individual bundles a single helix was taken from the Talin1 model and cloned multiple times to build up the structure. A short amino acid sequence (6 amino acids) was used to represent the region between helices and between bundles. The models were loaded into UCSF Chimera^67^ and a surface to show the protein-exterior boundary was added. The model was then exported as a .x3d file which can be imported into the animation software Blender (www.blender.org). Talin was assembled and an armature added to the mesh. This allows the structure to be easily “posed” in different forms while maintaining the correct size. The vinculin structure was unfolded first in PyMOL using the rotate function to ensure that protein size is respected. Then a similar process of surface-generation and armature-addition in Blender was followed.

### Assumptions

For the long unstructured regions, we used the mean contour length of an unstructured polypeptide, taken as 0.38 nm per amino acid in all cases^37^. For extended polypeptides the average width was taken to be 0.46 nm which is the approximate mean side chain width of a surface representation of a polypeptide.

### Bioinformatics

To determine protein similarity, we used the NCBI blastp tool available at https://blast.ncbi.nlm.nih.gov/. The sequences of *Homo sapiens* Talin1 (Q9Y490), *Homo sapiens* Talin2 (Q9Y4G6), *Caenorhabditis elegans* Talin (G5EGK1), *Dictyostelium discoideum* TalinA (P0CE95), and *Danio rerio* Talin1 (Q5U7N6) were gathered from UniProt (code in parentheses).

## Acknowledgements

BTG is funded by Biotechnology and Biological Sciences Research Council (Grant BB/S007245/1) and Cancer Research UK Program Grant (CRUK-A21671). SFHB is funded by the Max Planck Guest Scholarship program.

We thank Katy Goult, Ian Goult, Jen Hiscock, Neil Ball, Tom Paige, Vaclav Veverka, Melissa Andrews, Yulia Revina and Ada Cavalcanti-Adam for feedback and reading of the manuscript.

## Supplementary information

### Supplementary Figures

**Supplementary Figure 1:**
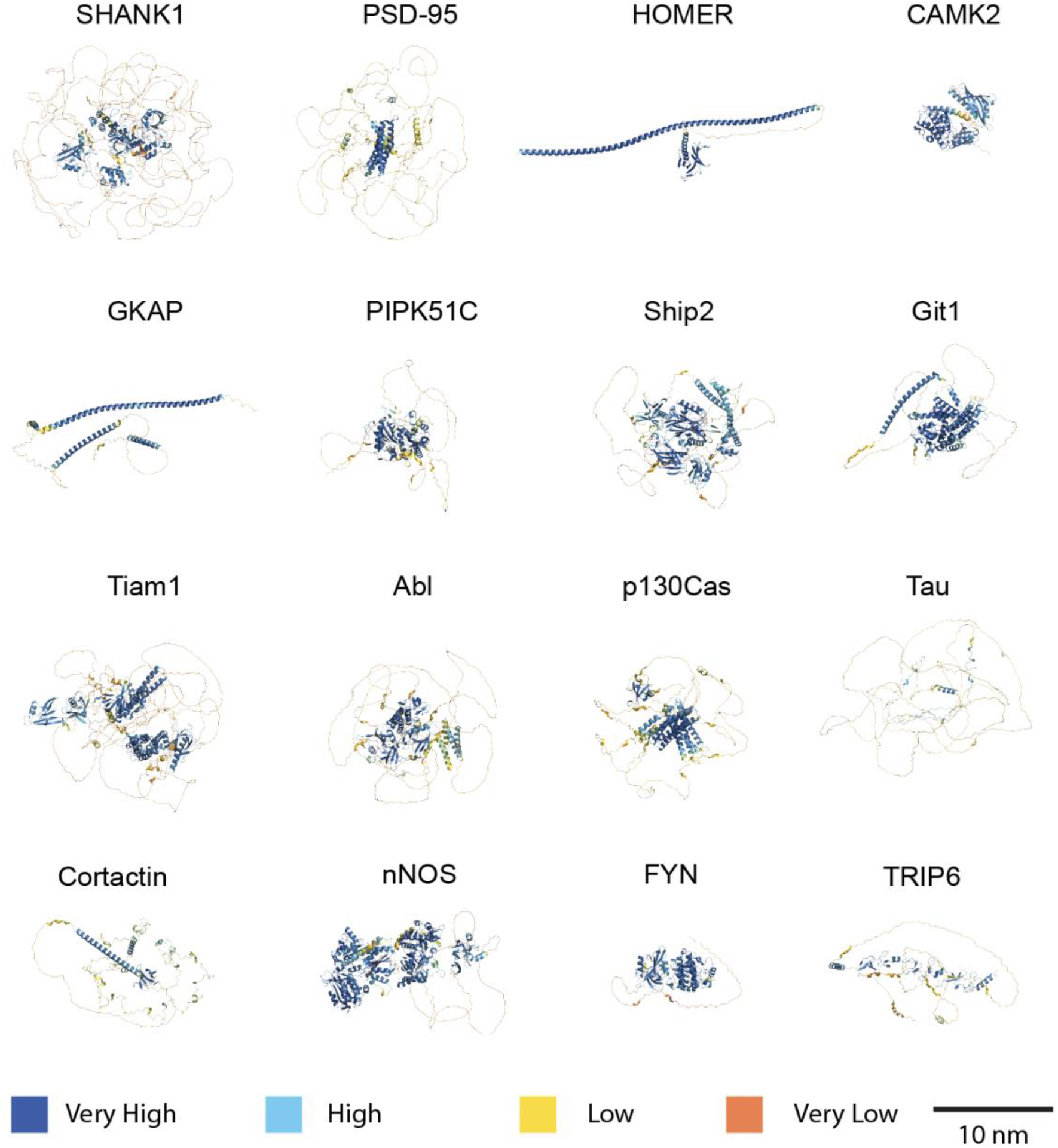
Selection of synaptic proteins and Integrin Adhesion Complex proteins with large expanses of unstructured regions generated using AlphaFold. The colour coding indicate how often amino acids end up with the same neighbours. Dark blue indicates stable domains. Orange indicates unstructured regions.

**Supplementary Figure 2:**
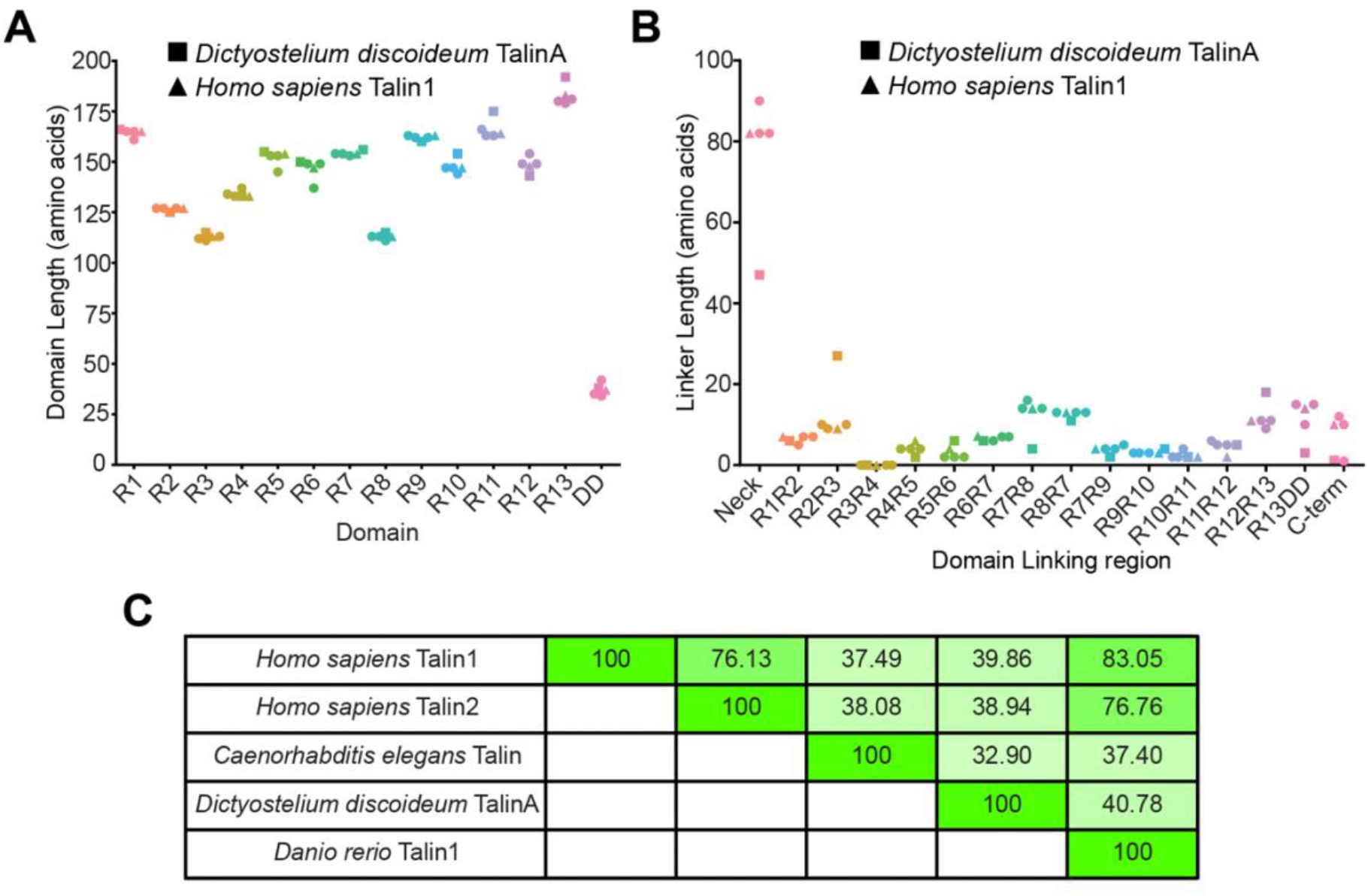
Talin protein domain structure is evolutionarily conserved. **A)** Measurement of domain sizes of talin proteins from different species that have diverged over the last 0.6-1 billion years with *Homo sapiens* Talin1 and *Dictyostelium discoideum* TalinA shown to highlight this conservation. Other proteins are *Homo sapiens* Talin2, *Caenorhabditis elegans* Talin, and *Danio rerio* (zebrafish) Talin1. **B)** Measurement of the amino acid sequence that bridge the talin domains in A. e.g. R1R2 denotes the linker region between the R1 and R2 domains. C-term represents the amount of unstructured sequence following the dimerization domain. **C)** Similarity matrix showing the sequence conservation between the proteins in A-B.

### Supplementary Movies

**Supplementary Movie 1. Talin R3 as the paradigm of a force-dependent binary switch**. All 13 talin switches are shown in the folded, 0 state. Mechanical force opens the R3 switch, causing it to switch binding partners between RIAM (yellow) bound to R3(0) and two vinculin (purple) bound to the R3(1) states. https://youtu.be/1R1CO9cOoRE

**Supplementary Movie 2. Talin undergoes dramatic alterations in length**. Three talin molecules are shown in different states, all engaged to an integrin (blue) at the N-terminus. Folded – the 13 rod domains are all in the 0 state. Unfolded – the 13 rod domains are all in the 1 state in the helical form. The positioning of the 11 vinculin binding sites (red helices) is shown. Stretched – the 13 rod domains are all in the fully extended 1 state where it is extended to the completely unstructured polypeptide chain form. The length of the talin molecule ranges from 86 nm (folded) to ∼350 nm (unfolded) to 800 nm (stretched). The diameter of a dendritic spine is ∼1000 nm. https://youtu.be/yCOkSaIkAQ4

**Supplementary Movie 3. MeshCODE – binary information written into the shape of a talin molecule**. A single talin molecule and the corresponding binary string is shown. At start of animation all the switches are in the folded, 0 state. As force is exerted on the molecule, the switches open and close independently and the binary string is updated. N.B. Only the string of helices, 1 state is shown and not the fully extended 1 state, this is for visualisation purposes, in reality each helix in the open 1 state would extend from 5 nm to 12 nm. https://youtu.be/pqVANarhQi4

**Supplementary Movie 4. A zone of activity of a protein tethered to the talin R8(0) switch**. The molecule has a tether at one end and an enzymatic domain at the other end. Attached via its tether site, the enzymatic domain can reach anywhere within the sphere but cannot reach outside of that zone. The radius of the sphere is a physical property of the linker region of the enzyme but the location of the tether where that sphere is attached is a function of the talin switch patterns. https://youtu.be/90WZYos01LU

**Supplementary Movie 5. A single posttranslational modification (PTM) dramatically alters the MeshCODE dimensions**. A movie showing how a chemical modification to the talin switches can alter the 3-dimensional location of molecules within the cell. A single talin molecule is shown attached to integrin (blue). A zone of influence of a molecule on the R9 switch is shown (grey sphere) positioned 87 nm away from the integrin attachment. The enzyme cyclin dependent kinase 1 (CDK1) is shown interacting with the R8(0) switch, and phosphorylating talin. This phosphorylation destabilises R7R8, and the two domains unfold, introducing a 110 nm extension to the talin molecule. As a result the zone of influence of the molecule on R9 is now 110 nm further from the integrin attachment. https://youtu.be/LVpwCPePovk

